# Distance-dependent distribution thresholding in probabilistic tractography

**DOI:** 10.1101/2022.07.27.501671

**Authors:** Ya-Ning Chang, Ajay D. Halai, Matthew A. Lambon Ralph

## Abstract

Tractography is widely used in human studies of connectivity with respect to every brain region, function, and is explored developmentally, in adulthood, aging, and in disease. However, the core issue of how to systematically threshold, taking into account the inherent differences in connectivity values for different track lengths, and to do this in a comparable way across studies has not been solved. The study adopted Monte Carlo derived distance-dependent distributions (DDDs) to generate distance-dependent thresholds with various levels of alpha for connections of varying lengths. As a test case, we applied the DDD approach to generate a language connectome. The resulting connectome showed expected short- and long-distance structural connectivity in the close and distant regions within the language network. The finding demonstrates that the DDD approach can be used for both individual and group thresholding. Critically, it offers a standard method that can be applied to various probabilistic tracking datasets.

---

Recent methodological advances in diffusion neuroimaging and probabilistic tractography have enabled the delineation of white matter fibre pathways *in vivo* [1-3]. For example, studies have reconstructed pathways in the human brain that are known to sustain higher-level cognitive functions such as language where non-human analogues are difficult to define and track [4-10]. Whilst probabilistic tractography allows us to investigate fibre pathways in vivo, there are at least two fundamental challenges with interpreting outputs. First, there is no consensus on how to threshold connectivity matrices, which impacts on the ability to perform statistical analyses that are comparable across studies. Second, brain networks typically comprise of short- and long-range connections but the latter inherently have distance dependant reductions in streamline likelihoods due to the inherent nature of probabilistic tractography. To gain a better understanding of long connections and networks, a more sophisticated treatment for distance effects is required. Thus, the present study set out to tackle these two key issues.

While the probabilistic tracking approach has proved useful in reconstructing complex structural connectivity networks, it has critical challenges especially with respect to false positives [11-14]. By definition, probabilistic tractography is designed to explore all possible streamlines, which are generated from fibres orientation distributions with differing degrees of non-random noise. Consequently, this method often produces widely connected brain networks [14]; however, thresholds can be applied to allow for more interpretable outcomes and is particularly useful in reconstructing complex crossing fibres [12]. Previous research has introduced several thresholding approaches. At the individual level, one could apply a set threshold to all individuals to remove sub-threshold connections or apply different thresholds across individuals to retain the same number of connections [15-17]. Group-level networks are commonly generated based on the consensus of connections, which retains connections present in some fraction of participants (i.e., at least 50% of subjects) [16, 18]. However, typically the threshold is arbitrary and as such a range of thresholds are often used to characterise the network [17]. Taking the field as a whole, given that there are inconsistencies in methodologies across datasets, tracking procedures and algorithms, it makes formal comparisons across studies difficult.

Regardless of the specific threshold used, it is common to apply a uniform threshold to all connections regardless of length/distance. This could be problematic, especially in a large-scale distributed brain network, because noise accumulates and streamline likelihoods decrease as a function of path length [19]. Thus, if a strict threshold is applied in order to minimise false positives near the seed region, then it is likely that long-range connections would not exceed that threshold, resulting in distance false negatives. A recent study by Roberts et al. [14] developed consistency-based thresholding, which computed variation of unthresholded connectivity strength across individuals. A threshold was applied to connections with high consistency (i.e., low variation) across the group to preserve a desired level of connectivity density. It was assumed that if long-range connections were truly connected, the connections would be consistent across individuals such that consistency could be high even though the connection strengths were low. Given that the individual connectivity matrix was not thresholded, it remains unclear if this approach may overestimate connections as an unthresholded matrix is almost always fully connected [14]. Indeed, the authors noted that the consistency-based approach could potentially be biased by specific types of data acquisition and pre-processing. More recently, Betzel et al. [16] developed a distance-dependent consensus thresholding approach. This approach is similar to the consistency-based approach in that no threshold was applied at the individual level while group-level consensus was obtained by computing the fraction of individuals that had connection weights greater than zero. The novel aspect is related to applying a correction based on distance (across a range of bins), where for each bin the pair of brain regions that have the highest consensus across individuals is preserved. One potential caveat is that the bins used should be reasonably wide to keep the same number of pairs as in the original matrix. Also, similar to Roberts et al. [14], this approach was primarily operated at the group level.

Despite the fact that a variety of thresholding approaches have been proposed, critical issues remain. There is still a lack of empirical support for a common thresholding regime that allows for the comparison across studies and one that resolves the distance artefact at both individual and group levels. A previous study by Cloutman et al. [7] did propose an approach towards standardisation that was inspired by statistical hypothesis testing. In that study, probabilistic tracking was conducted to map connectivity between the sub-regions of the insula and the rest of the brain. For each individual, the distribution of connection scores between the seed region and each region of an atlas was fitted with a Poisson distribution, and that was used to identify a threshold value at α = 5%. At the group level, the consensus approach was used, in which connections were selected only when they were consistently identified across at least 50% or 75% of participants. This distribution-thresholding approach at the individual level was the first attempt to determine individual thresholds based on a statistical metric (alpha) using the connectivity distributions; however, it did not take into account distance effects, as in Betzel et al. [16] and Roberts et al. [14]. The present study extended this distribution approach by developing a distance-dependent distribution (DDD) method in order to establish a common ground for thresholding based on significance levels of alpha. Specifically, as per standard Monte Carlo type statistical methods, we generated normative distributions of randomly-sampled connectivity and then set different levels of alpha to identify the corresponding thresholds. Critically, we generated the DDDs at different distances to derive thresholds for connections of varying length. To illustrate the novel approach, we applied DDD thresholding to generate a language connectivity matrix and evaluated if it could capture *a priori* language networks reported in the literature [4, 5, 8, 9]. Specifically, it was expected that the resulting connectivity matrix could capture long-range connections and reproduce key white matter tracts in the language network including arcuate fasciculus (AF) and/or superior longitudinal fasciculus (SLF), which are generally associated with the dorsal language pathway; and middle longitudinal fasciculus (MdLF), inferior frontal-occipital fasciculus (IFOF) and uncinate fasciculus (UF), which are generally associated with the ventral language pathway.

## Results

Figure 1A shows the result of the tractogram from an axial slice for a representative participant in diffusion space. The colour indicates directionality; red for right-left, green for anterior-posterior and blue for superior-inferior. Figure 1B shows an example of the extracted streamlines connecting two ROIs in the inferior frontal cortex. We extracted streamlines between all of the ROIs pairs to compute connectivity strength. Figure 1C shows the 260 ROIs covering the left hemisphere only.

**Figure 1.**
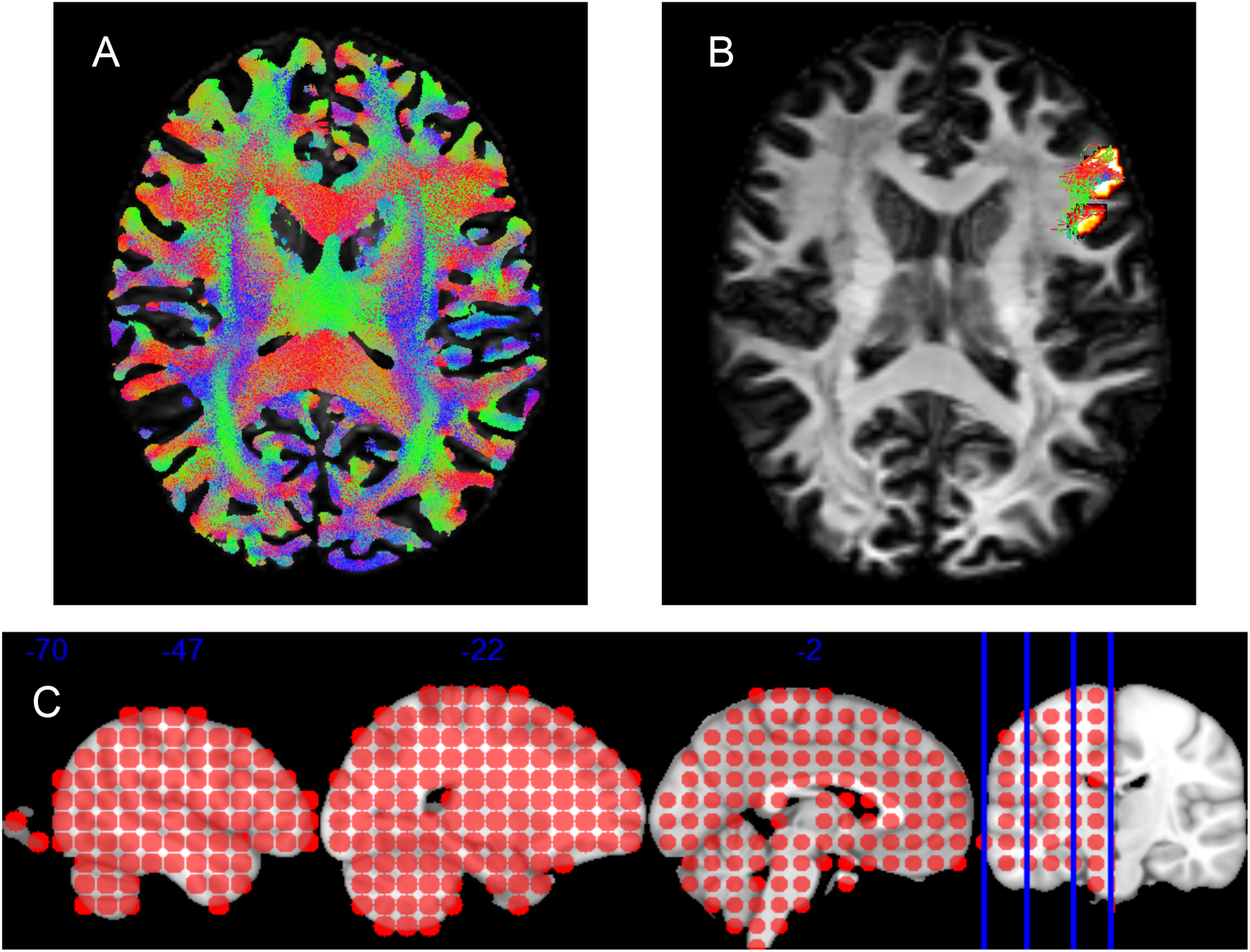
(A) The representative whole-brain tracking with 10M streamlines. (B) An example of the extracted streamlines connecting two ROIs in the inferior frontal cortex. (C) The 260 ROIs (diameter 8mm) covering the left hemisphere.

### Distance-dependent distributions (DDDs) and thresholds

Figure 2 shows the number of ROI paired samples for 26 distance groupings from the 86 unique distances. For each distance range, there were at least 1000 samples, ranging from 1068 to 2844. Note that the range was not equally spaced due to the limitation of retaining a minimum number of samples per bin; however, we explored a different distance categorisation, and the key results remain similar (see Supplementary, Figures S1-S4).

**Figure 2.**
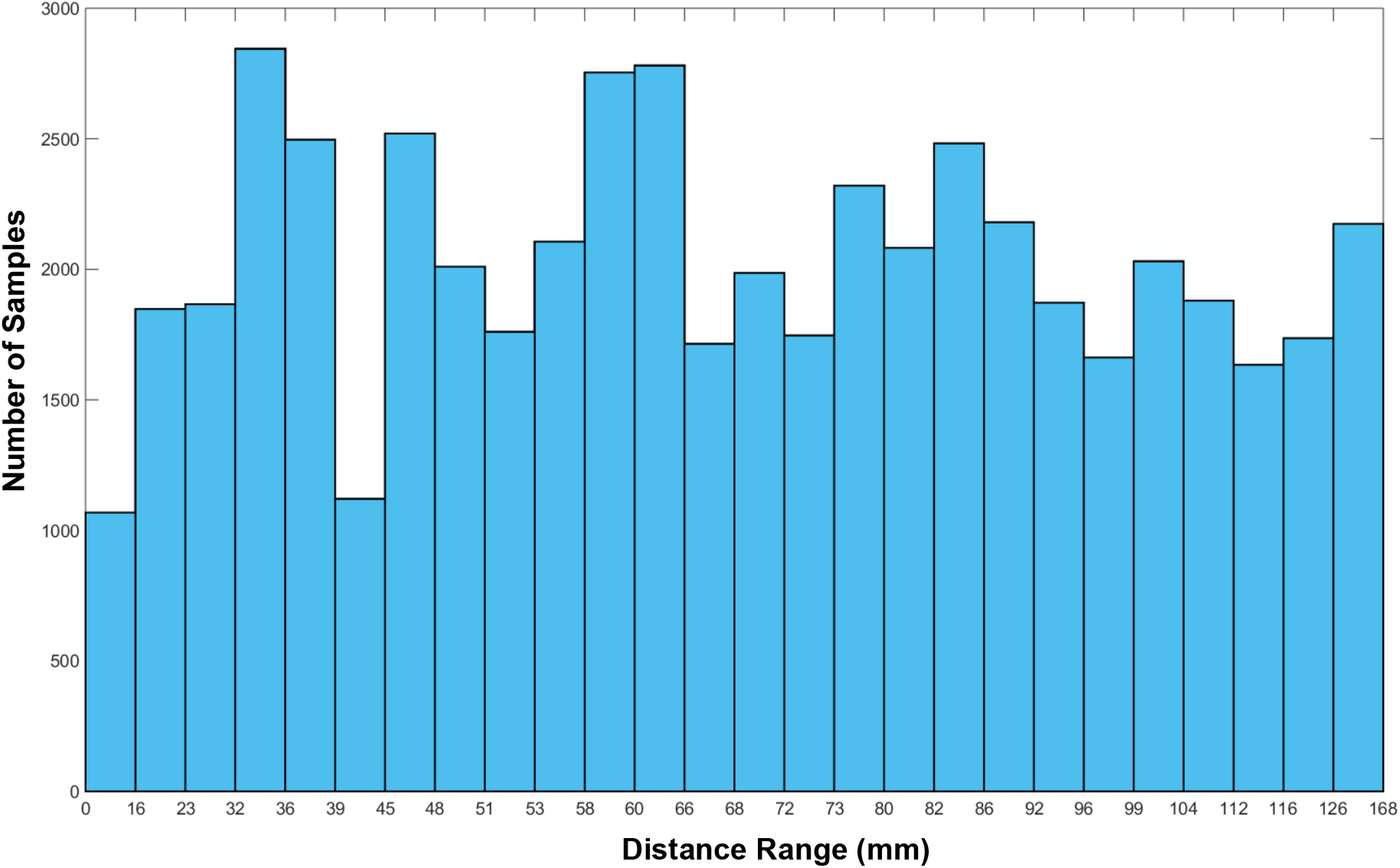
The number of ROI paired samples for each of the 26 distance ranges.

For each distance range, we first averaged the connectivity scores across participants and then conducted resampling 100,000 times to generate a null distribution of random connectivity. Figure 3 shows the sampling distributions for each distance range. As expected, the short-range connections tended to have higher connection scores compared to the long-range connections, as noted by the right skewed distributions. For the longer connections, the majority of connection scores were close to zero; however, as expected given the presence of long-range white matter and fasciculi in the brain, there were notable extremes that had strong connectivity differed from the majority of the null connectivity.

**Figure 3.**
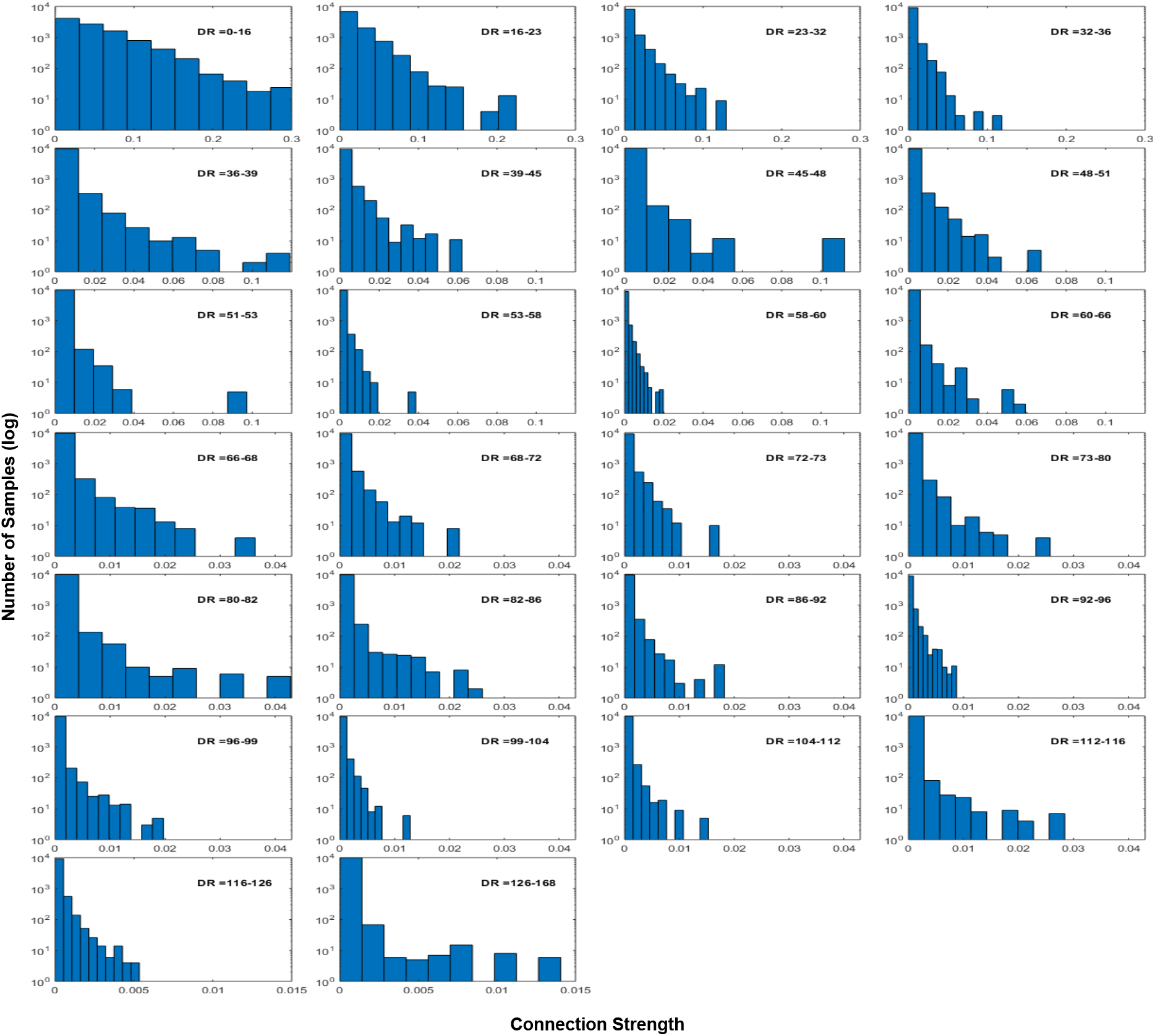
The sampling distribution of random connectivity for each of the 26 distance ranges. The x-axis indicates connection strength and the y-axis indicates the number of samples on a log scale. DR: distance range (mm).

Figure 4 shows that the DDD thresholds at three alpha levels of 10%, 20% and 30% across the 26 distance bins. Specifically, for each alpha level, the connectivity threshold generally decreased with increased distance. As a result, the short-distances had higher thresholds than long-distance connections. Moreover, the connectivity thresholds were moderated by the alpha level, where the highest thresholds were noted for the smallest alpha and the lowest thresholds for the largest alpha. Next, we applied the DDD thresholds to generate a language connectivity matrix.

**Figure 4.**
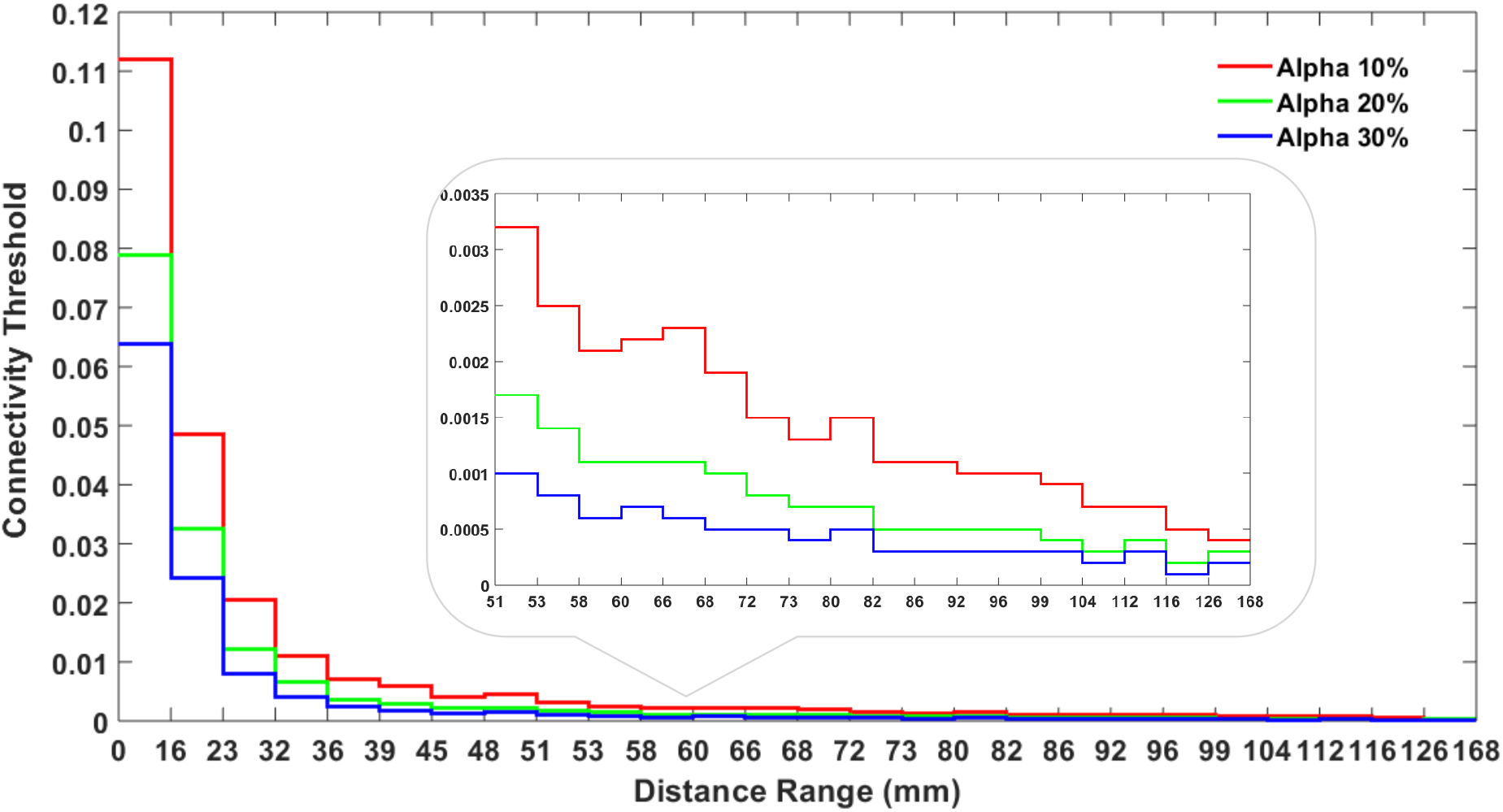
The distance-dependent distribution thresholds at three alpha levels of 10%, 20% and 30% varied with the 26 ROI distance bins.

### Language network connectivity

Table 1 shows the distance between 13 language ROIs included in the analyses (See Methods for details). The distance varied widely as expected, where IFG Tri and pFG were furthest apart (101.45 mm) and the mFG and pFG were closest together (16.28 mm). We first converted all distance values to integers, which were used to compute the average language connectivity matrix across individuals. The three levels of the DDD thresholds were applied to the outputs (as in Figure 4) according to the distance of the ROI pairs. Note that we did not threshold the connectivity matrix at the individual level because the DDD approach did not require sparse individual connectivity matrices which are typically required for group-level consensus thresholding. For completeness, we also demonstrated that the DDD thresholds could also be applied to individual connectivity matrices, where the thresholded individual matrices had high correlations with the thresholded average matrix (see Supplementary Figures S5-S6).

**Table 1.**
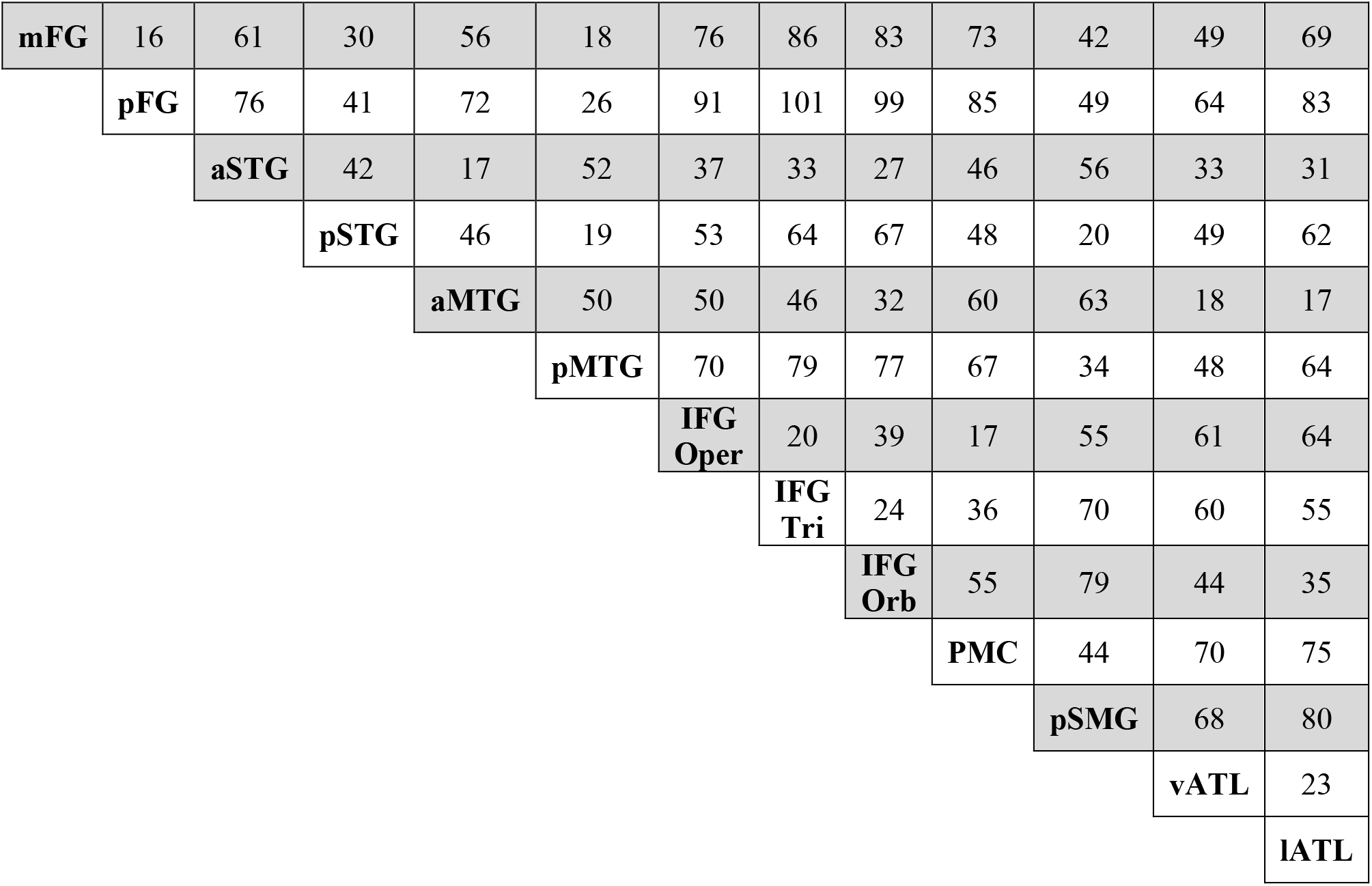
The distance (mm) between all language ROIs.

After thresholding, the average connectivity score was binarised and coded with three different colours according to the alpha levels (red = 10% of alpha, green = 20% and blue = 30%) as in Figure 5.

**Figure 5.**
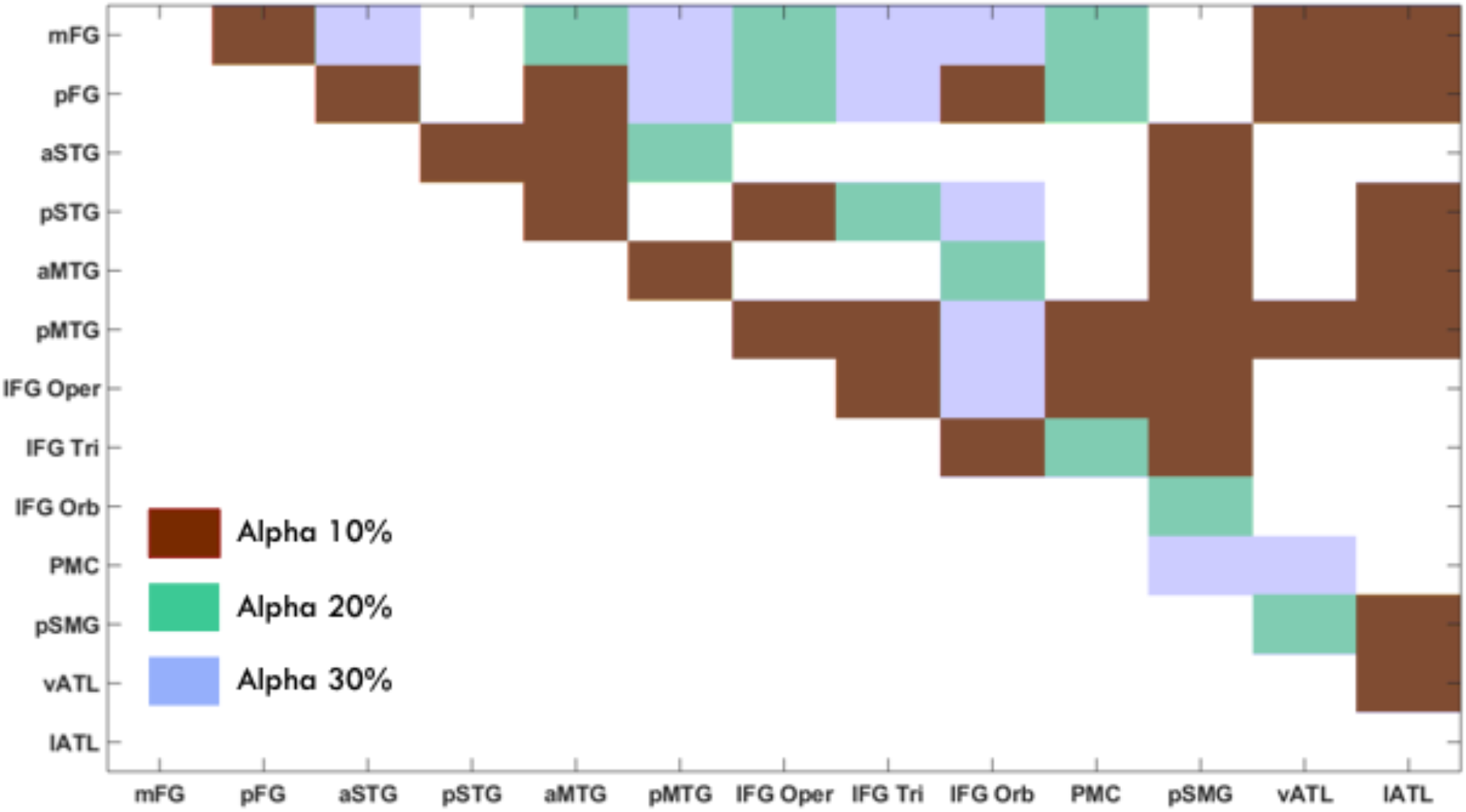
The average language connectivity matrix across individuals after thresholding and binarisation. The different levels of alpha with 10% on the top, 20% on the middle, and 30% on the bottom were superimposed onto one matrix wherein anything that survives the 10% would also survive 20% and 30%.

We projected the connection scores (without binarisation) to the brain for an intuitive way to visualise the results (Figure 6) using the BrainNet Viewer [20]. The width of the line indicates the strength of connection. With the most stringent threshold (i.e., the alpha level of 10%), the connectivity between the ROIs in both the close and distant cortical regions was reconstructed. Specifically, we observed neighbouring ROIs to be connected in the occipito-temporal regions (mFG and the pFG); temporal lobe (aSTG with pSTG and aMTG with pMTG); anterior temporal lobe (vATL and lATL); and frontal lobe (IFG Orb, IFG Tri and IFG Oper) as well as premotor cortex (PMC). We also observed distant ROIs being connected between the frontal and the parietal lobes, the temporal and the parietal lobes, the frontal and the temporal lobes, and the frontal and the occipito-temporal lobes. Specifically, the IFG Tri and IFG Oper were connected to the pSMG presumably via the AF, which could be part of the dorsal language pathway. The aSTG, pSTG, aMTG and pMTG were also connected to the pSMG presumably via the MdLF, which could be associated with the ventral language pathway. Moreover, there was long-range connectivity between the IFG Tri and IFG Oper and the pMTG via the AF, as well as between the IFG Orb and the pFG via the IFOF and/or part of the ILF, which could be part of the ventral language pathway.

**Figure 6.**
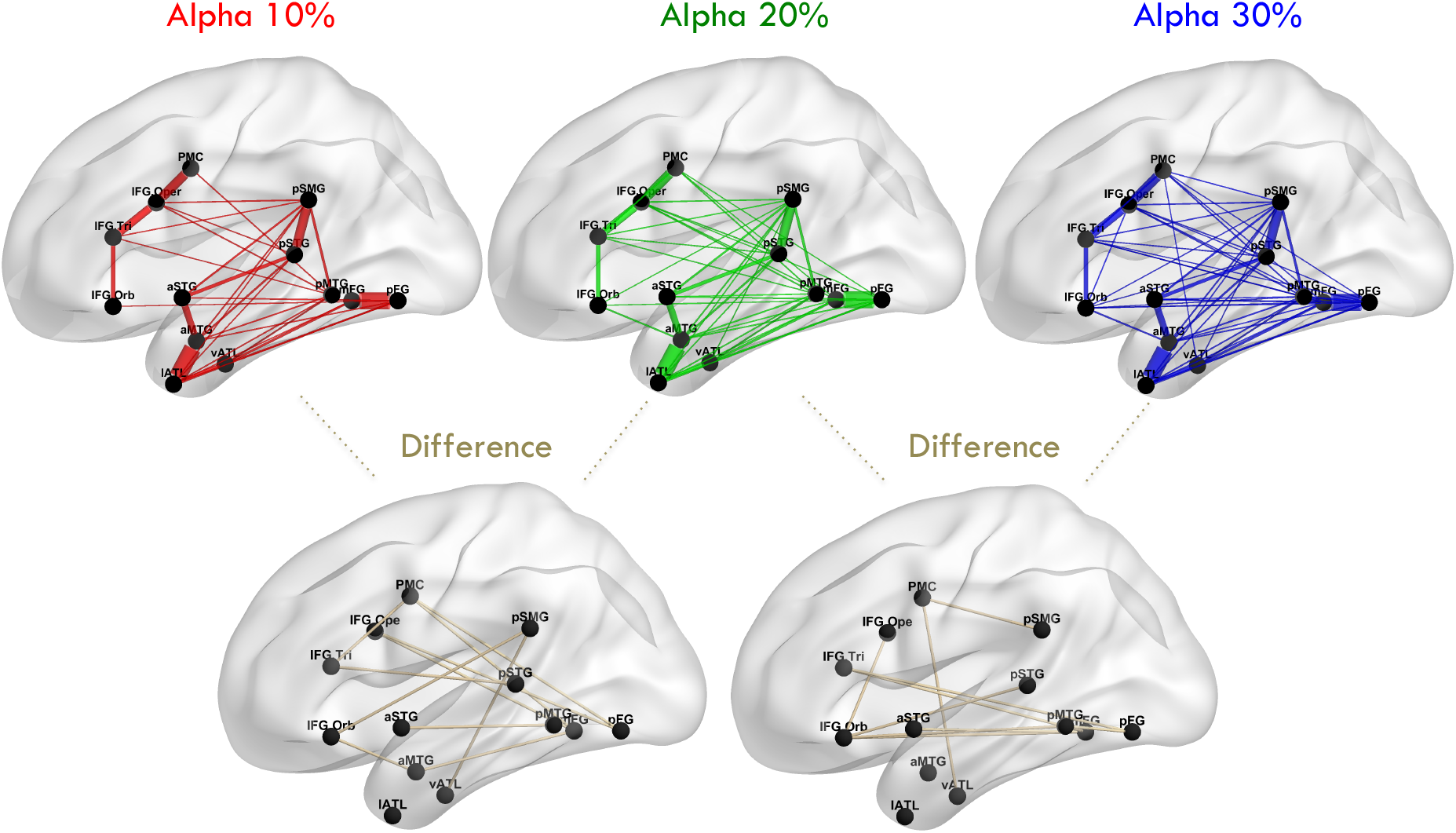
The projections of the average language connectivity matrix in the brain with three alpha levels of 10% (red), 20% (green) and 30% (blue) and the difference plots between them (gold). The wider the connection line indicates the stronger the connections.

With lenient thresholds (i.e., the alpha levels of 20% and 30%), more cross-region connectivity could be observed, especially for the inferior frontal, temporal and occipital regions. The IFG Orb was connected with the aMTG via the uncinate fasciculus (UF), and the IFG Oper and the IFG Tri were connected to the mFG and pFG via the AF. Collectively, these results demonstrated that the connectivity matrix based on the DDD thresholding approach was able to reconstruct the key white matter tracts that sustain language processing in the language network as previously reported in tractography studies [4, 5, 8, 9] and the three inferior frontal regions had different connectivity profiles with temporal and parietal regions, consistent with the cortico-cortical evoked potential study [21].

## Discussion

Probabilistic tractography has become an increasingly important tool in neuroimaging studies to delineate white matter fibre pathways *in vivo*. The key advantage of probabilistic tractography is that it can reveal all possible tracts from a seed because each streamline propagates along a direction drawn from a probabilistic distribution. The main limitation, however, is unavoidably increasing the possibility of producing false connections [11-14]. Thus, the primary aim of this study was to develop a thresholding approach for probabilistic tractography that is grounded in statistical hypothesis testing and to resolve the false negatives commonly associated with long-distance connections. Specifically, we used a data-driven distance-dependant distribution (DDD) approach to generate normative distributions of random connectivity for 26 distance ranges based on ROIs spanning the left hemisphere. We demonstrated how this novel method could be applied to uncover the connectivity profile for a language network and how different alpha levels affect the final outcome. Critically, we were able to reproduce the expected language network [4, 5, 8, 9] as shown in Figure 6. Overall, this study demonstrated that the data-driven and statistical-inspired thresholds can minimise false positives for short-range connections and false negatives for long-range connections.

To date, most connectivity thresholds are chosen heuristically and, arguably, arbitrarily based on existing literature or exploratory outcome. For a given dataset, one could use a more lenient criterion to explore the probable tracts in contrast with a more stringent criterion, which could be used to identify core bundles. Indeed, it has been suggested that networks might be better characterised with a broad range of thresholds [17]. Additionally, thresholding might also be related to the demographic, psychosocial and medical information of the individuals, such as age, gender and mental illness [22]. Although there may be justifications to apply specific thresholds to a given dataset, the outcomes may not be comparable with or generalise to other datasets. The ability to compare across studies is critically important in validating and testing the reliability of key findings. Thus, while studies have focused on probabilistic tractography and improving tractography algorithms [23-26], only a handful of studies have developed thresholding approaches that make the resulting connectivity close to the ‘ground true’ or comparable across the species [14, 16, 18]. Our data-driven DDD approach was designed to work with any probabilistic tracking dataset and to provide a common ground (i.e., the alpha level) to relate thresholds across datasets regardless of the specific tracking approach and parameters used (provided that random sampling distributions are established per dataset). Obviously, if studies have used similar datasets, tracking tools and parameters, then the same DDDs can be directly applied.

Our DDD approach dealt with the distance artefact by generating higher thresholds for close regions and lower thresholds for distant regions (Figure 4). As long-range connections tend to have smaller connection strength, some tools such as FSL [24, 26] apply a distance correction by multiplying the distance with the probability of connection strength. This can lead to probabilities greater than one, which makes interpretations difficult. More importantly, it is not clear what mathematical form best characterises the relationship between distance and the probability of connection strength for correction. Two studies have tried to overcome these challenges. Roberts et al. [14] introduced consistency-based thresholding, which does not directly deal with distance but it could effectively preserve long-range connections using high consistency thresholds (possible if variance across individuals is small). In contrast, Betzel et al. [16] developed distinct thresholds for the group-level consensus scores of different lengths. Although both approaches have proved their effectiveness for group-level thresholding, the approaches cannot be applied to individual-level thresholding. In contrast, we have demonstrated that our DDD approach can work directly with the average connectivity matrix to identify plausible connections related to language processing (Figure 5) and can also be applied at the individual level (see Supplementary). Thus, our DDD approach extends previous distance-related thresholding approaches, while supporting both individual-level and group-level analyses.

There are some limitations that are not addressed with the DDD approach. First, the DDDs generated by the present study may not be directly applied to tractography datasets that are acquired using different data acquisition protocols, pre-processing, tractography algorithms, or different numbers/sizes of ROIs. That is because the distributions of random connectivity are data-specific and generated from the outcome of a specific tractography setting. This means that DDDs would need to be calculated for each study/analysis; however, we have provided detailed methodological descriptions about how to generate appropriate DDDs (downloadable from https://www.mrc-cbu.cam.ac.uk/bibliography/opendata/). Despite the computational cost required to generate data-specific DDDs, a major advance is that one could formally compare results between studies based on the alpha threshold and in the future collated data for formal meta-analyses. Secondly, the ROIs used to generate DDDs cover the left hemisphere but not the right hemisphere. This was deliberate for two reasons: (a) crossing fibres through the corpus callosum are complex especially for the callosal fibres connecting from the midsagittal slice of the corpus callosum to inferior and lateral brain regions [27, 28]; and (b) to keep computational costs down. Further work is required to extend the principles outlined for the DDD approach to the right hemisphere and the whole brain. Lastly, our DDD approach is designed to work with the outcome of tractography and to provide a method of comparing thresholding results regardless of tractography algorithm/settings. Thus, we did not seek to identify optimal thresholds for a ‘ground truth’. Instead, we considered a range of comparable thresholds that can help characterise the structural connectivity [17] and bridge studies using different tractography approaches. We acknowledge the key role of thresholding techniques in reducing the probability of spurious connectivity in probabilistic tractography, which can allow us to identify ‘real’ networks.

To conclude, the DDD approach was developed as a strategy to formally threshold structural connectivity maps from any tractography datasets and allow for comparisons across studies. The DDD approach also addressed the distance artefact by providing different thresholds for short- and long-range connections.

## Methods

### Human connectome data, pre-processing and tracking

Fifty-four participants’ pre-processed structural and diffusion datasets were downloaded from the WU-Minn 1200 Human Connectome Project (HCP) [29-33]. Both the HCP T1-weighted and diffusion imaging data were acquired on a 3T Siemens Skyra “Connectome” scanner with a customised SC72 gradient insert and a customised body transmitter coil with 56 cm bore size.

The HCP T1-weighted (T1w) images were acquired using the 3D Magnetisation Prepared Rapid Acquisition GRE (MP-RAGE) method with TR = 2400 ms, TE = 2.14 ms, T1 = 1000 ms, flip angle = 8°, FOV = 224 × 224 mm, voxel size = 0.7 mm isotropic, BW = 210 Hz/Px and acquisition time = 7 m 40 s. The diffusion weighted images were acquired using a Spin-echo EPI with TR = 5520 ms, TE = 89.5 ms, flip angle = 78 deg, refocusing flip angle = 160 deg, FOV = 210 × 180 mm, voxel size = 1.25 mm isotropic, and BW = 1488 Hz/Px.

In the HCP, a full diffusion session included six runs for three different gradient tables, and each table acquired twice, one from right-to-left and the other one from left-to-right phase encoding polarities. Each gradient table included approximately 90 diffusion weighting directions plus six b = 0 acquisitions interspersed throughout each run. Multiple shells of b = 1000, 2000 and 3000 s/mm^2^ were utilised with each shell having an approximately equal number of acquisitions within each run.

The dataset used in the present study were pre-processed using the HCP pipelines. Briefly, the T1w and T2w images were aligned in native space using FSL’s FLIRT and FNIRT functions. A field map distortion correction was conducted and registered to T1w and T2w images using FSL’s FLIRT boundary-based registration (BBR) algorithm. The structural images were subjected to bias field correction [30]. Each participant’s native structural volume space was registered to MNI space using FSL’s FLIRT and FNIRT functions. As the diffusion data were collected with reversed phase encoded polarities, these pairs of images were utilised to estimate the susceptibility-induced off-resonance field, and they were combined into a single image using FSL’s TOPUP and EDDY functions for distortion-correction [34-36].

The distortion-corrected data were submitted to the MRtrix3 toolbox (https://www.mrtrix.org/) for further processing and whole-brain probabilistic tracking. Specifically, the diffusion data were first subjected to bias correction using the ANTS flag [37]. Next, a response function was estimated using spherical deconvolution for grey, white matter and cerebrospinal fluid (CSF) compartments using the ‘dhollander’ algorithm.

Subsequently, we averaged the response function across subjects for each tissue type to obtain a group average, which was then used to estimate fibre orientation distributions (FOD) using the multi-shell multi-tissue constrained spherical deconvolution algorithm [38]. Finally, intensity normalisation (in the log-domain) was applied to all FOD outputs. Whole-brain tractography was performed using MRtrix3 with anatomically constrained priors (obtained using the five-tissue segmentation function) and the iFOD2 algorithm. We obtained 10 million streamlines per subject with a maximum streamline length of 250 mm and a fractional anisotropy (FA) cut-off value of 0.06. The resultant whole-brain tractogram was further filtered using spherical-deconvolution informed filtering of tractograms (SIFT2) to improve the quantification and biologically-meaningful nature of whole-brain connectivity [23].

### Distance-dependent distributions of random connectivity

To generate normative distributions via randomly-sampled connectivity at different distances, we created a grid of 230 regions of interest (ROIs) with a diameter of 8 mm that covered the entire left hemisphere in MNI space. For each participant, ROIs were inverted to native diffusion space (using inverse FLIRT and FNIRT transforms) and binarised over the grey and white matter interface (GWI). The GWI-ROIs were used as a mask to extract the number of streamlines connecting all other ROIs based on the whole brain tractogram. The connection value between a pair of ROIs was quantified by computing the proportion of streamlines emerging from the seed and ending at the target ROI. It was expected that closer ROI pairs would have higher connection scores while distant ROI pairs would have lower scores. However, as the ROIs were randomly placed, many ROI pairs would have connection scores close to zero.

The pairwise connection scores between ROI were collapsed and the number of samples for each ROI distance was computed. For simplicity, we computed the distance between two ROI coordinates and rounded it to the nearest integer, resulting in 87 unique distances (range = 16 -168 mm). As there were varying numbers of samples for each distance, the distance values were further grouped into 26 distance bins to ensure that each had at least 1000 samples. For each distance bin, the average connection values across participants was computed and randomly sampled 100,000 times to generate sampling distributions. The procedure of resampling generated 26 distance-dependent distributions of random connectivity. For each distribution, we set three alpha levels of 10%, 20% and 30%, which resulted in three different DDD thresholds.

### Language network ROIs

To generate a language connectivity matrix, we selected 13 left hemisphere ROIs (8 mm diameter) associated with language processing and reading. The peak coordinates were taken from the literature with slight modifications to avoid any overlap between ROIs [9, 39-43]. The ROIs included the middle and posterior fusiform gyrus (mFG and pFG), the anterior and posterior superior temporal gyrus (aSTG and pSTG), the anterior and posterior middle temporal gyrus (aMTG and pMTG), the opercularis, triangularis and orbital parts of the inferior frontal gyrus (IFG Oper, IFG Tri and IFG Orb), the premotor cortex (PMC), posterior supramarginal gyrus (pSMG) and both the lateral and ventral anterior temporal lobe (vATL and lATL). The language connectivity matrix was generated by computing the connection strength between all of the language ROIs pairs. A group-level language connectivity matrix was generated by computing the average connectivity matrix across individuals. We then applied the three levels of the DDD thresholds to the average language connectivity matrix according to the distances between the ROI pairs.

## Data Availability

The image data used in this work was downloaded from the WU-Minn 1200 Human Connectome Project (http://www.humanconnectomeproject.org/).

## Code Availability

The relevant scripts to generate DDDs are available (conducted in MATLAB) from the MRC Cognition and Brain Science Units Data Repository (https://www.mrc-cbu.cam.ac.uk/bibliography/opendata/).

## Acknowledgements

This research was supported by grants from the European Research Council (GAP: 670428 - BRAIN2MIND_NEUROCOMP to MALR), Medical Research Council intramural funding (MC_UU_00005/18 to MALR), the Rosetrees Trust (A1699 to ADH and MALR) and the Ministry of Science and Technology (MOST110-2423-H-006-001-MY3 to YC).

