## Supplementary for "Distance-dependent distribution thresholding in probabilistic tractography"

### SUPPLEMENTARY MATERIALS

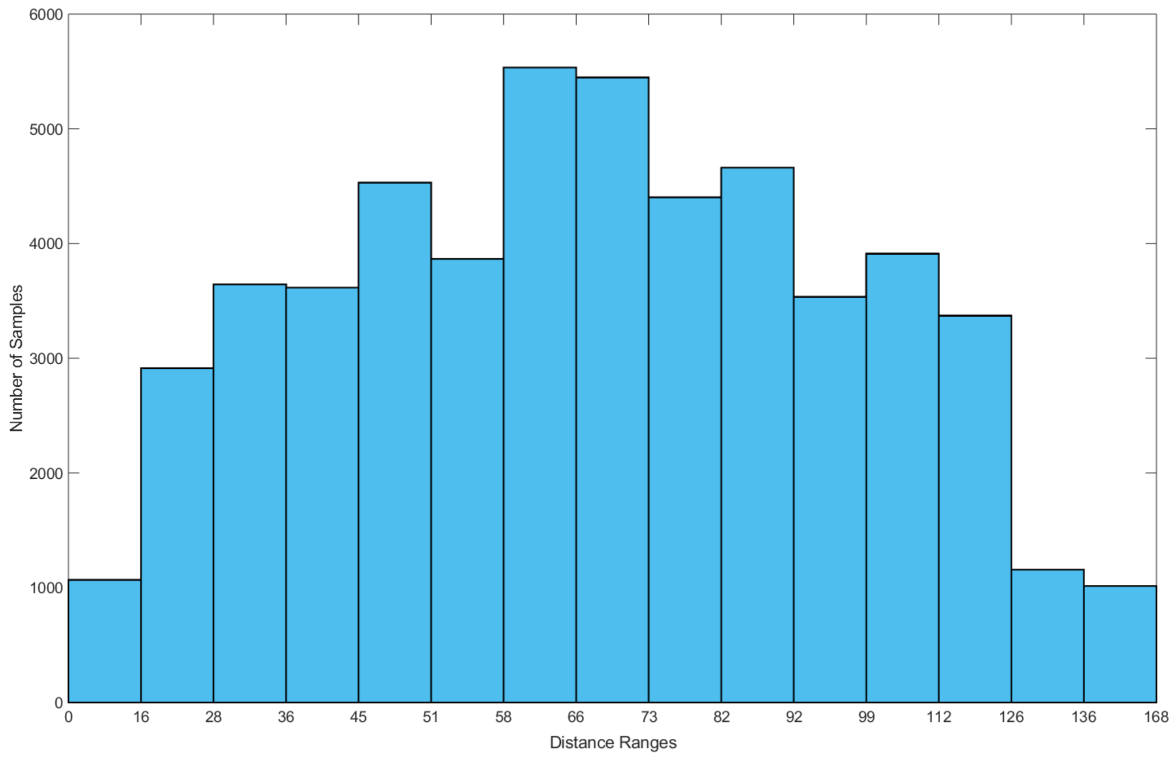


Figure S1. The number of the ROI paired samples categorised into 15 distance ranges, which is much less than the distance ranges reported in the main text.

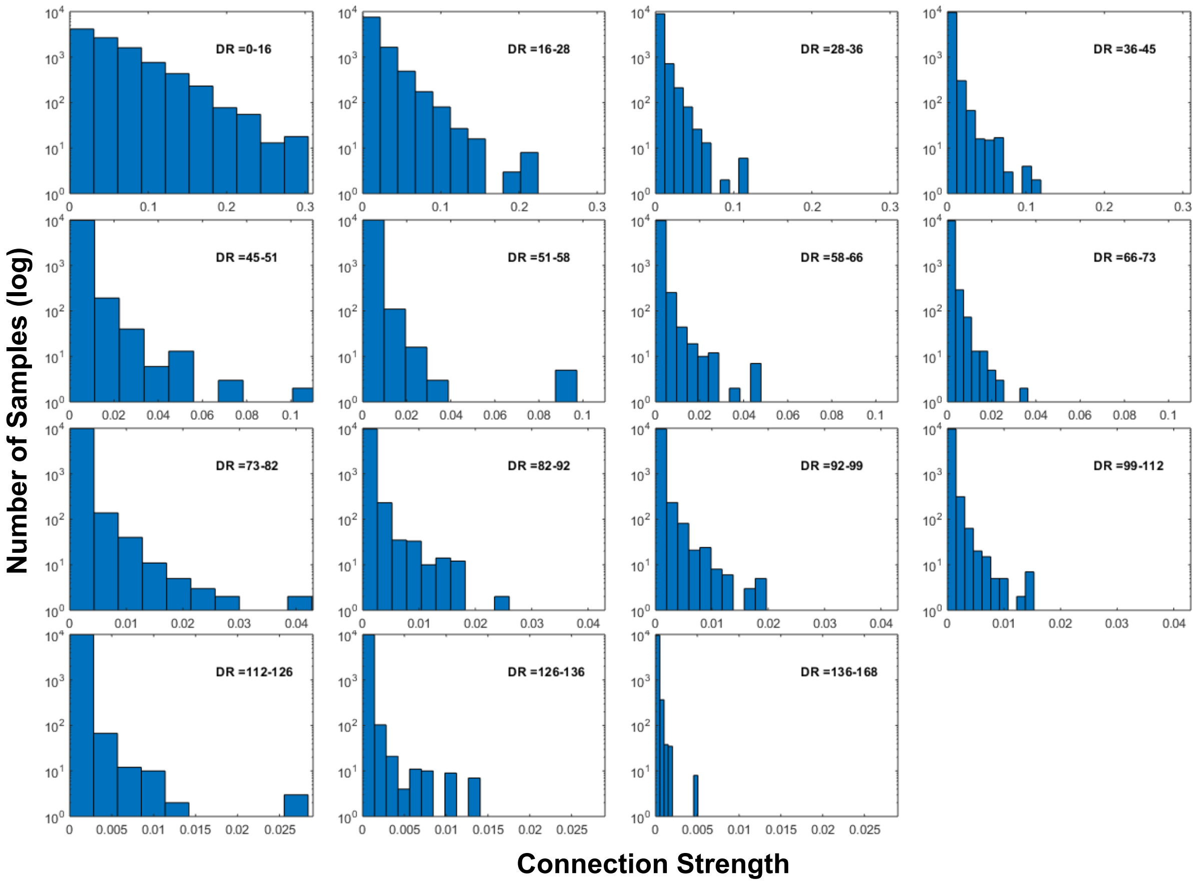


Figure S2. The sampling distribution of random connectivity for each of the 15 distance ranges. The x-axis indicates connection strength and the y-axis indicates the number of samples.

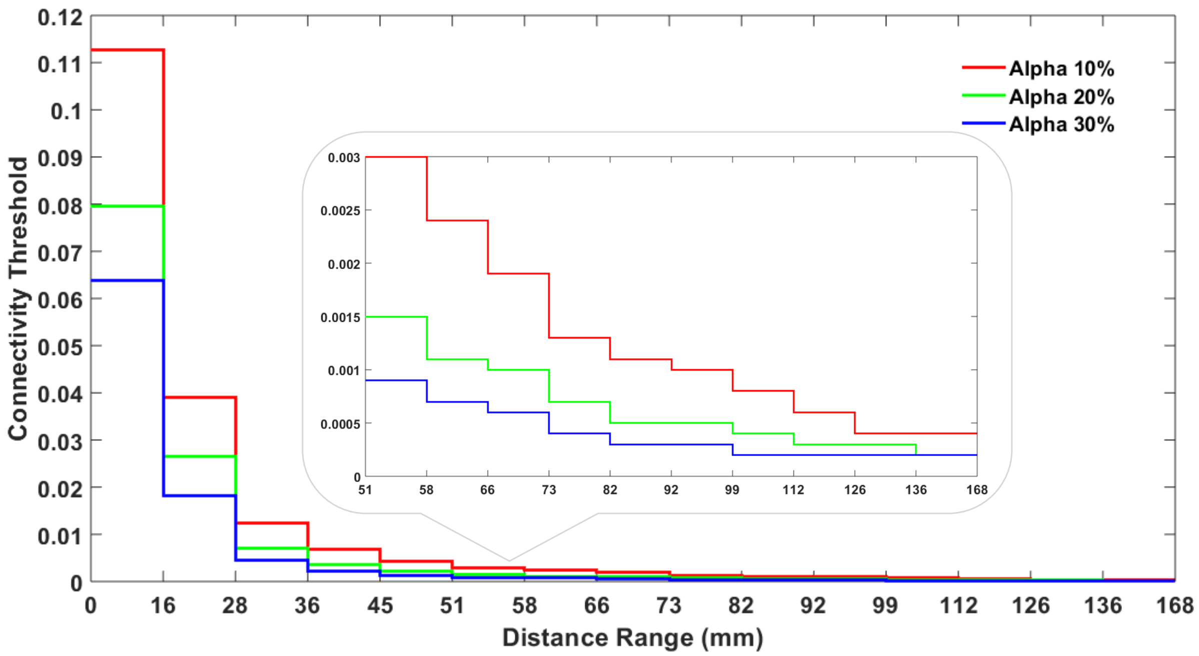


Figure S3. The distance-dependent distribution thresholds at three alpha levels of 10%, 20% and 30% varied with the 15 ROI distance ranges.

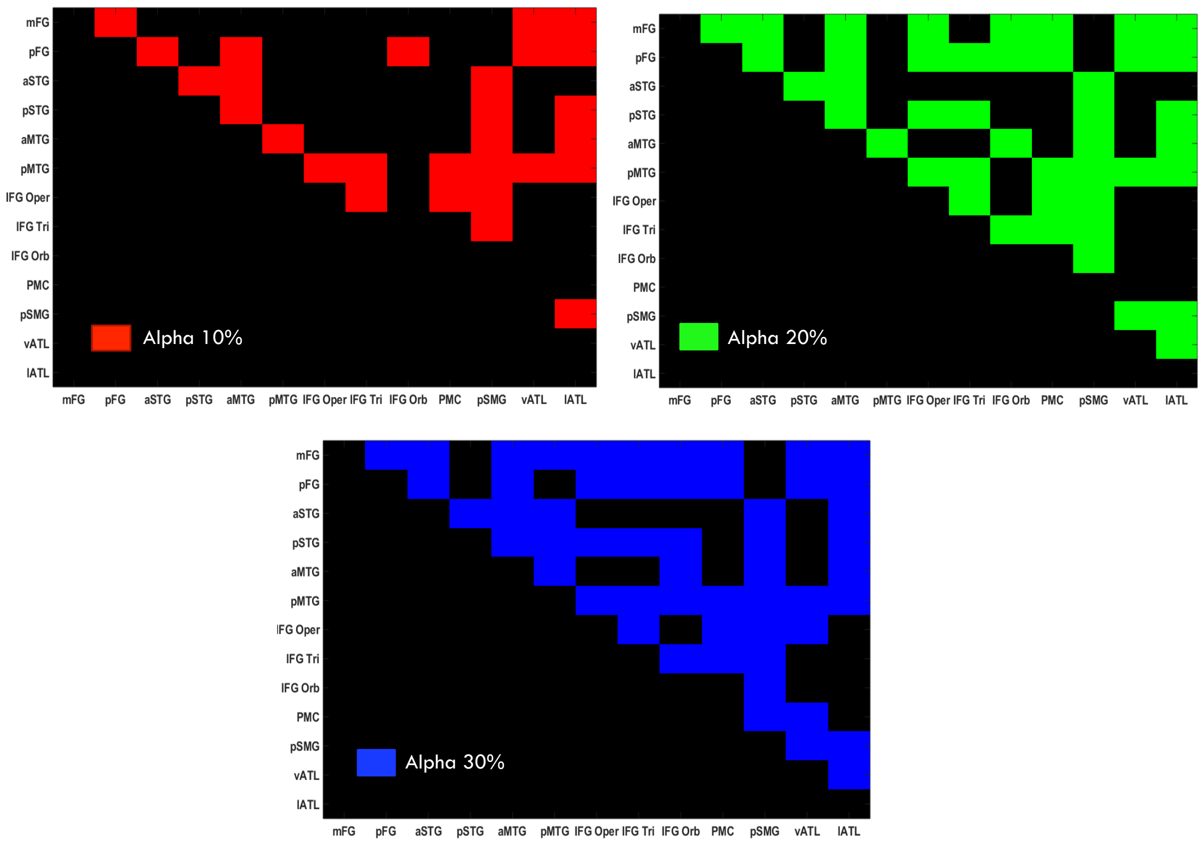


Figure S4. The average language connectivity matrix after thresholding based on the 15 distance ranges at the alpha levels of 10%, 20% and 30%. The connectivity matrix is very similar to Figure 5 reported in the main text. The dice similarity between the two matrices is 0.953.

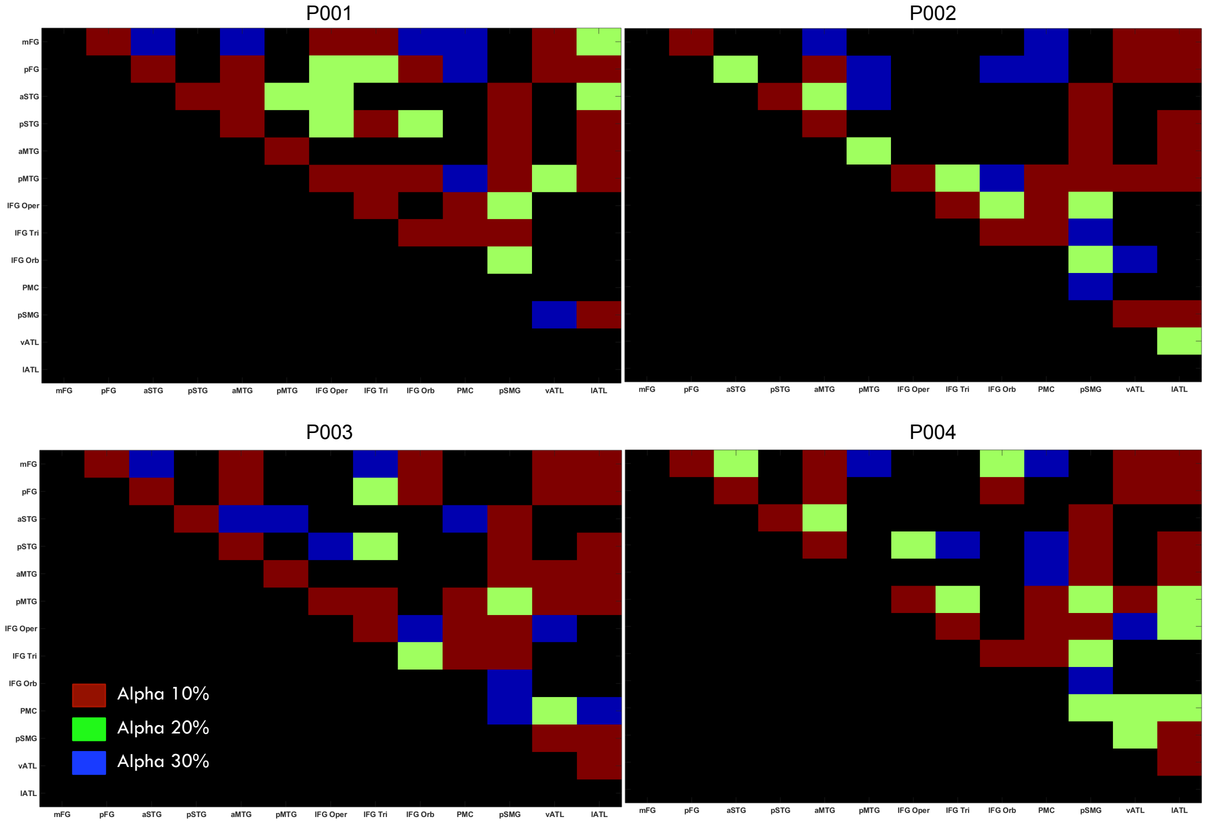


Figure S5. The individual language connectivity matrices for four representative participants after thresholding based on the 26 distance ranges at the alpha levels of 10%, 20% and 30%, as reported in the main text. Note that the colour scheme is hierarchical for simplification such that the green cells indicate thresholded connectivity in addition to the red cells, and the blue cells indicate thresholded connectivity in addition to both the red and green cells. As can be seen, there are individual differences in terms of brain connectivity and the strength of connectivity; however, the general pattern of connectivity is largely similar.

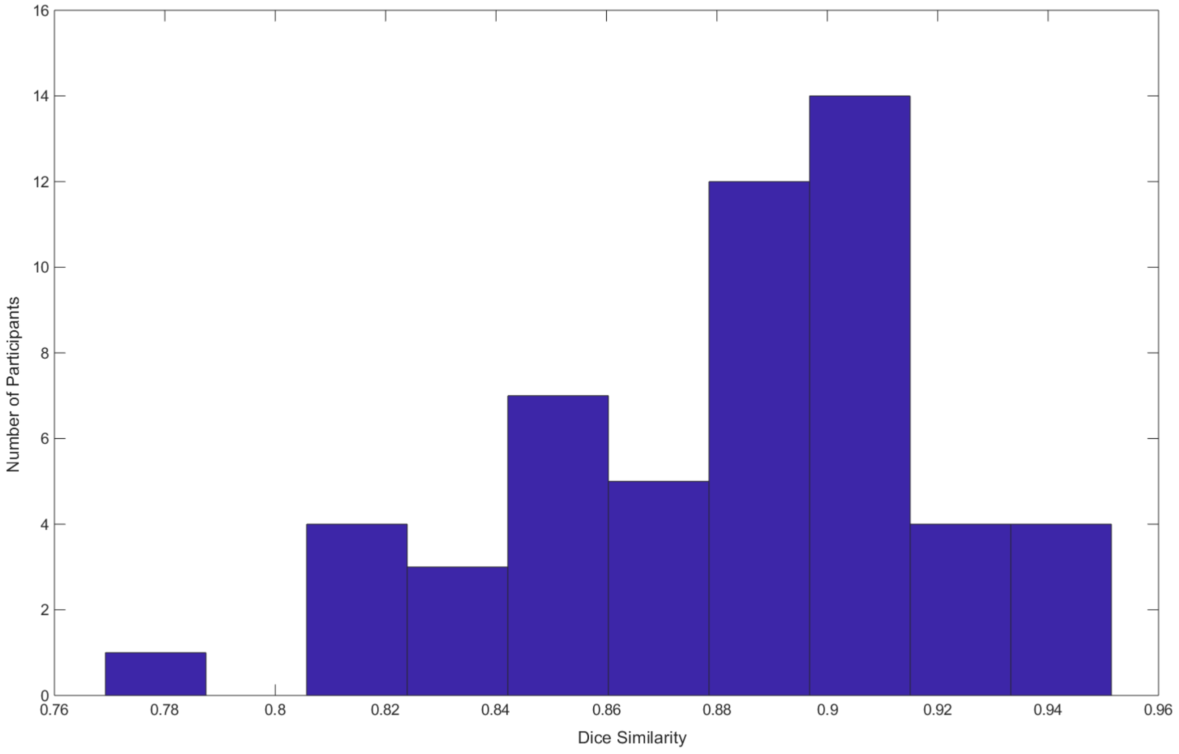


Figure S6. The dice similarity between the individual language connectivity matrices and the average language connectivity matrix for all of the 54 individuals. As can be seen, the similarity scores are generally very high (M=0.883, SD=0.038). The result suggests that the DDD thresholds can also be directly applied to the individual level.
